# Sugar feeding by invasive mosquito species on ornamental and wild plants

**DOI:** 10.1101/2023.04.13.536683

**Authors:** Irving Forde Upshur, Mikhyle Fehlman, Vansh Parikh, Chloé Lahondère

**Author notes:** To whom all correspondence should be addressed: Chloé Lahondère Department of Biochemistry; Virginia Polytechnic Institute and State University Blacksburg, VA, 24061, USA.

## Abstract

Feeding on plant-derived sugars is an essential component of mosquito biology that affects key aspects of their lives such as survival, metabolism, and reproduction. Mosquitoes locate plants to feed on using olfactory and visual cues. *Aedes aegypti* and *Aedes albopictus* are two invasive mosquito species to the US, and are vectors of diseases such as dengue fever, chikungunya, and Zika. These species live in heavily populated, urban areas, where they have a high accessibility to human hosts as well as to plants in backyards and town landscapes. Therefore, it is important to understand what plants may attract or repel mosquitoes to inform citizens and municipal authorities accordingly. Here, we analyzed *Ae. aegypti* and *Ae. albopictus* sugar-feeding behavior with eleven different commonly planted ornamental plant species. We then assessed feeding activity using the anthrone method and identified volatile composition of plant headspace using gas-chromatography mass-spectroscopy. Finally, we determined the sugar-feeding activity of field caught mosquitoes using the plant DNA barcoding technique and compared these results with the eleven ornamental species tested in the laboratory. The potential for application to disease vector control is also discussed.

## 1. Introduction

Phytophagy, the act of feeding on plants, is important for many insect species, including blood-sucking mosquitoes. Acquiring carbohydrates is essential for both males and females and can, in some species, constitute the sole source of food for adults (*e.g.*, *Toxorhynchites* spp.). Males feed exclusively on plant-derived sugars and recently emerged females tend to seek for sugar before taking their first blood-meal, which enhances egg production [1]. In addition to carbohydrates, it has been determined that mosquitoes also acquire vitamins, amino acids, and salts from plant nectar [2–4].

To locate a sugar meal, mosquitoes are driven by several cues, including visual and olfactory (*e.g.*, plant-emitted semiochemicals) [5]. Volatile odorant compounds (VOCs) are perceived *via* odorant receptors present primarily on the antennae, which are fine-tuned to specific volatiles and elicit attractive or repellent responses [6,7]. Terpenes and benzenoids are common compound classes in flower scent profiles which drive mosquito attraction. Some mosquito species have been shown to be attracted to flower semiochemicals like linalool, (Z)-3-hexen-1-ol, and benzaldehyde and repelled by compounds such as β-myrcene and limonene [5,8]. A more comprehensive understanding of which plant-emitted semiochemicals are attractive and repellent to mosquitoes is critical to the development of new and efficient disease vector control tools. This is a pressing undertaking, as current control efforts are being challenged by increased mosquito insecticide resistance worldwide. [9].

A promising mosquito control strategy is the use of attractive toxic sugar baits (ATSBs) containing mixtures of attractive odorants and toxic compounds that take advantage of mosquitoes’ natural requirement to feed on sugar. These traps bypass pre-existing problems that conventional control strategies have faced, such as insecticide resistance, and have been shown to have a low impact on non-target organisms like plant pollinators and mosquito predators [10,11]. In ATSBs, a chosen toxin (*e.g.,* boric acid, dinotefuran) is mixed with sugar in water and supplemented with an organic volatile attractant (*e.g.,* eugenol, garlic oil), after which the mixture can be applied to a plant, fruit, or bait-station [12,13]. Several studies have identified volatile compounds from fruits and plants that mosquitoes locate and feed on in the field [14,15]. However, these volatiles are usually mosquito species-specific and, within the wide range of olfactory stimuli mosquitoes experience in the field, it remains unclear what compounds are the most attractive. Successful ATSB use has been observed in many different mosquito species including *Aedes aegypti*, *Aedes albopictus*, *Aedes japonicus*, *Culex pipiens*, *Culex quinquefasciatus*, and *Anopheles gambiae* [11,16–19].

Among current disease vectors that are of particular concern are *Ae. aegypti* and *Ae. albopictus* mosquitoes. These species are responsible for spreading dengue, chikungunya and Zika viruses, all of which have frequent global incidence as vaccines and / or treatments remain unavailable [20]. According to a recent study by Leta *et al.* (2018), a total of 215 countries and territories exhibit environments that are suitable for *Ae. aegypti* and *Ae. albopictus* habitation. In the context of climate change and global warming, the geographic distribution of these mosquitoes might widen and potentially spread diseases in new areas, making it crucial to develop control strategies for both species [21].

Sugar feeding in mosquitoes is relatively understudied compared to host-seeking and blood-feeding behavior, as pathogens are transmitted to humans and animals when infected female mosquitoes bite. Yet it appears crucial to study this behavior, as it bears the potential for the development of new tools for vector surveillance and control. Resources that mosquitoes use in populated urban areas to obtain a sugar meal and how invasive species adapt to local ornamental plants remain poorly understood. Some ornamental plants (*e.g.*, *Ligustrum quihoui*, *Pittosporum tobira*, *Loropetalum chinense*) have been shown to increase survivorship in *Ae. albopictus* and could therefore contribute to their ability to successfully transmit disease by increasing their overall fitness [22]. In addition, the distribution of a population of *An. gambiae* male mosquitoes in Western Burkina Faso was shown to be influenced by the presence or absence of ornamental plants [23]. Moreover, ornamental plant abundance has been shown to directly affect the population distributions of both *Ae. aegypti* and *Ae albopictus* during field experiments in Huixtla, Chiapas for *Ae. aegypti* and Guangzhou, China and Long Island, New York for *Ae. albopictus* [24–26]. When tested for the presence of sugar in their crop, individuals from both species exhibited a higher proportion of sugar feeding when collected from an urban area with a higher amount of blooming ornamental plants, compared to urban areas with little or no ornamental plant presence. This prompts the hypothesis that certain ornamental species, by providing nectar to the mosquitoes, might greatly influence their fitness and could consequently increase the risk of disease transmission in heavily populated areas.

The present study aims to provide a better understanding of sugar feeding preferences in *Ae. aegypti* and *Ae. albopictus* in the US. We first examined mosquito landing and feeding preference on different ornamental plant species that are commonly found in nurseries and backyards. These plant species varied in flower shape, size, color, scent, and nectar contents, so that the possible effect of flower morphology and sugar availability on preference could be observed. We then analyzed the scent profile of each of these plants using gas-chromatography coupled with mass-spectrometry (GC-MS) to identify chemicals that may attract or repel mosquitoes. Lastly, we used plant DNA barcoding on field caught mosquitoes to identify plants that they are feeding on. This work improves our understanding of the elusive mosquito-plant relationship and brings insights on the development of novel and ecologically friendly control tools such as attractive baits or repellents.

## 2. Materials and Methods

### 2.1. Insects

The Rockefeller *Ae. aegypti* strain (MRA-734, MR4, AATCC®, Manassas, VA, USA) and the ATM95 *Ae. albopictus* strain (ATM-NJ95, AATCC®, Keyport, NJ, USA) were used for this study. Larvae were reared in 26 x 35 x 4 cm covered trays that were filled with deionized water. The trays were kept in a climatic chamber at 26°±0.5°C and 60 ± 10% humidity under light:dark cycles of 12h:12h. The diet of the larvae consisted of Hikari Tropic First Bites (Petco, San Diego, CA, USA). Prior to starting the plant visitation experiment, around 100 pupae were placed into mosquito breeding containers (BioQuip, Rancho Dominguez, CA, USA—1425, 1425DG) on the day of pupation. Upon emergence, female and male mosquitoes were starved for 1-2 days before being individually selected with forceps. To do so, the containers were placed in a cool environment to immobilize mosquitoes and isolate them for the following plant visitation assays.

### 2.2 Plant Visitation Assays

#### Plant visitation protocol

Two BugDorm-1 insect cages (BugDorm, DP1000) were placed on top of a 26 x 35 x 4 cm tray containing DI water to help minimize risk of desiccation. These cages were then placed in a secondary larger acrylic cage (17 x 22 x 32 in) which further provided a warm and humid environment for the mosquitoes. Each assay was conducted with both mosquito species separately to allow for better comparisons between the two species (*e.g.*, survival, feeding). Within each cage, a GoPro camera (Hero5 Black), a water-containing cup covered by a wet paper towel for humidity, and the plant of interest were placed (Fig. 1A). Eleven ornamental plants were tested: wave petunia (*Petunia petunia x atkinsiana*), red impatiens (*Impatiens walleriana*), marigold (*Tagetes spp.*), *Celosia* (*spp.*), butterfly bush (*Buddleja spp.*), *Guara* (*spp.*), *Lantana* (*spp.*), Mexican heather (*Cuphea hyssopifolia*), *Scaevola* (*spp.*), goldenrod (*Solidago spp.*), and yarrow (*Achillea millefolium*). In addition, an iButton (Maxim, DS1923) was programmed and added to the cage to record humidity and temperature throughout the assays. Ten females and ten males of *Ae. aegypti* and *Ae. albopictus* were released into the cages at the end of the day between 4:30 and 5:30 pm, which has been previously reported as a peak sugar feeding activity time for *Ae. aegypti* and the beginning of the sugar feeding rhythm for *Ae. albopictus* [27,28]. In the following morning, between 8:30 and 9:30 am (total: 16 hours), alive mosquitoes were collected using a Bug Vacuum (Redeo, XCQ-B), sorted by sex, and stored for subsequent sugar analysis at −70 °C. Dead mosquitoes were tallied and removed before beginning another assay to assess survival. Three replicates, totaling thirty mosquitoes per sex per mosquito species, were conducted for each plant.

**Fig. 1.**
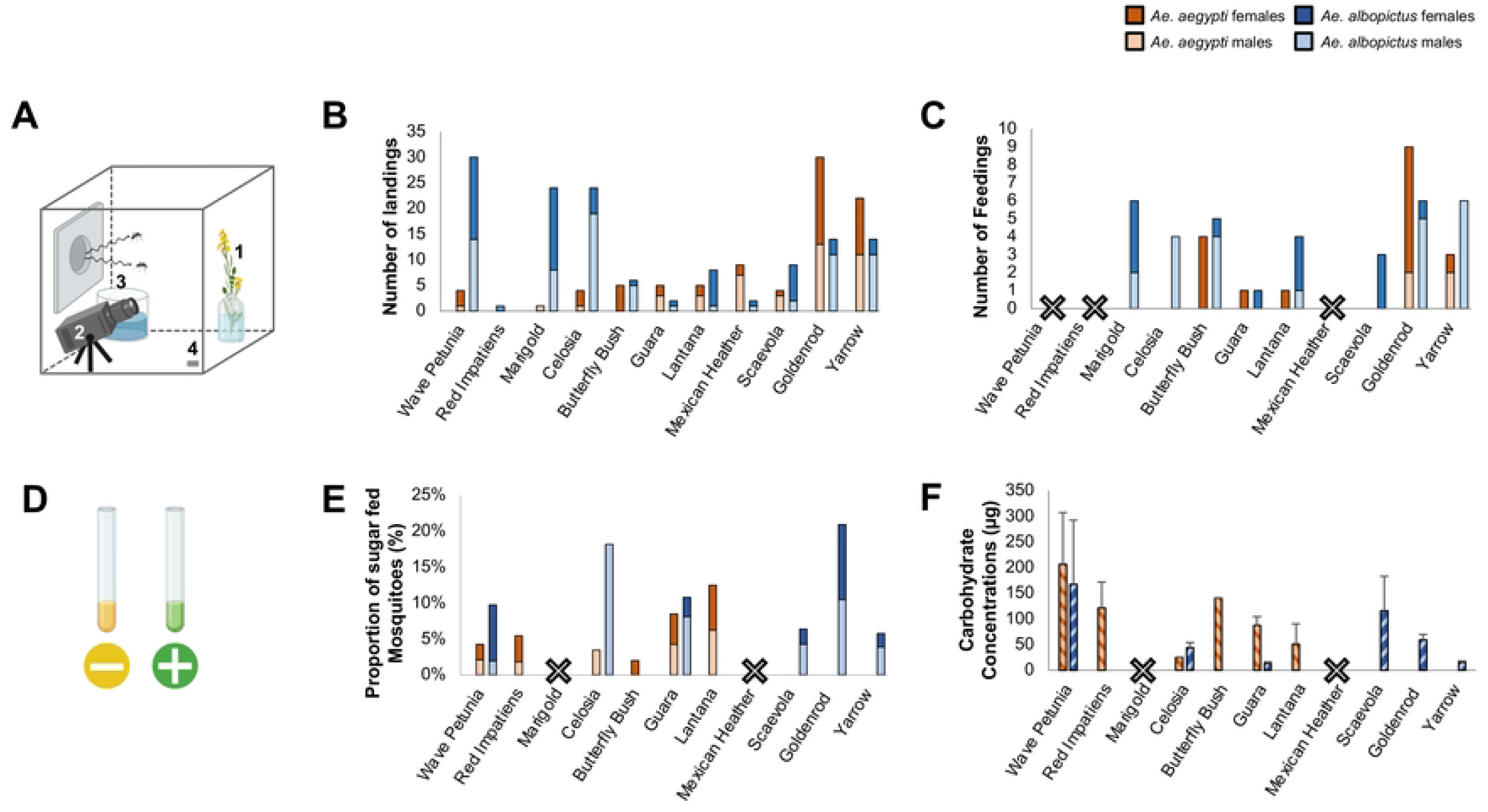
Mosquito sugar-feeding behavior on common ornamentals (A) The standard layout for a plant visitation assay. Mosquitoes are released in a cage containing the ornamental flower [1], a GoPro aimed at the flower [2], a water cup topped with a soaked paper towel [3], and an iButton temperature/humidity recording device [4]. (B) Total number of observed landings by either *Ae. aegypti* males (light orange) and females (dark orange) or *Ae. albopictus* males (light blue) and females (dark blue) on the eleven different tested ornamental flower species. (C) Total number of observed feedings. Ornamental flowers with a large ‘X’ indicate no activity from either species. (D) Negative (yellow) and positive (green) anthrone tests for the consumption of sugar. (E) Percentage of male and female *Ae. aegypti* and *Ae. albopictus* that tested positive for the consumption of fructose. (F) Total carbohydrate concentrations (µg) as determined by the anthrone method. Red impatiens and *Ae. albopictus* data is not present in these results (E, F), as an *Ae. albopictus* colony had not been established at the point of testing red impatiens.

#### Video analysis

GoPros cameras were used to record the mosquito behavior for a duration of two hours. Due to the consistent timing of initializing the assays and beginning the recording, we ensured that the videos captured the peak of sugar feeding activity in these day-active mosquitoes (between 4:30 p.m. and 7:30 p.m.). Each video was analyzed by counting the total number of mosquito landings and feedings on the flower of interest. Landings were defined as a mosquito flying to the flower and idling on it for any amount of time. A single feeding was defined as the vertical movement of a mosquito’s head in a flower (*i.e.*, probing), which is a characteristic motion of nectar-feeding. The sex of each mosquito landing or feeding was also noted for each landing and feeding event.

### 2.3 Scent Collection and GC-MS analyses

#### Scent collection

To collect the headspace of each plant species, the inflorescence of the plant was enclosed in a nylon oven bag (Reynolds Kitchens, USA) that was taped tight around the stem. Two tygon tubes (Cole-Parmer, USA) were connected on one side to a diaphragm air pump (Gast, Benton Harbor, MI, USA), while the other side was inserted at the small opening of the bag. Air flow was then initiated by connecting the pump to a 6V battery (Power-Sonic Batteries, USA). One tube pumped ingoing air into the bag through a charcoal filter cartridge (1 L/min), which served to remove any contaminants from the pump or the surrounding environment. The other tube pulled air out of the bag (1 L/min) through a headspace trap composed of a borosilicate Pasteur pipette (VWR, Radnor, PA, USA) containing 100 mg of Porapak powder Q 80-100 mesh (Waters Corporation, Milford, MA, USA). After 24 hours of headspace collection, traps were eluted with 600 μL of 99% purity hexane (Sigma Aldrich, 227064-2L). The samples were sealed in 2 mL amber borosilicate vials (VWR, Radnor, PA) with Teflon-lined caps (VWR, Radnor, PA) and were subsequently stored at −70°C to remain stable until analysis by GC-MS. For controls, samples were taken concurrently from empty oven bags. For each plant species, between 7 and 12 collection replicates were performed.

#### GC-MS analysis

An aliquot of 20 μL was pipetted from each plant scent sample from the preceding method and placed into a vial with an insert (VWR, Radnor, PA) to be analyzed with a GC-MS (Thermo Fisher Scientific, Trace 1310) equipped with a 30 m column (Thermo Fisher Scientific, I.D. 0.25 mm, #36096-1420). Helium was used as the carrier gas at a constant flow of 1 cc/min. After each sample was prepared for the species of interest, they were loaded into the machine using an autosampler (TriPlus RSH, Thermo Fisher Scientific). The oven temperature was set at 45°C, held for 4 minutes, followed by a heating gradient ramping to 230°C, held for 6 minutes (total run: 28.5 min.).

Chromatogram peaks were integrated using the Chromeleon software MS quantitative processing method (Thermo Fisher Scientific) and tentatively identified using the online NIST library. Major peaks found with consistently high abundances across multiple samples for each ornamental were then recorded for comparison across ornamental species. An internal standard, made of 100 ng/µL of heptyl acetate (Sigma-Aldrich, CAS #112-06-1), was added to each sample to calculate the concentrations of each compound based on a calibration curve. External synthetic standards, if available commercially, were used to confirm the chemical identity as in Lahondère *et al.* (2020).

### 2.4. Carbohydrate content assays

#### Nectar detection in the mosquito crop

Carbohydrate contents were measured by sequentially using the cold and warm anthrone methods described by van Handel (1985) [29]. First, mosquitoes were crushed with a glass rod in culture glass tubes (Sigma-Aldrich, C1048-72EA) containing 300 μL of cold anthrone reagent to detect for fructose consumption, as fructose is a monosaccharide that would only be present in the mosquito if it fed on a plant. The anthrone reagent was prepared by combining 150 mL water in a 1 L Erlenmeyer flask on ice with 380 mL sulfuric acid (Fisher, CAS #7664-93-9), in which 750 mg of anthrone (Sigma-Aldrich, CAS #90-44-8) was then dissolved. The samples were kept idle at room temperature (25°C) for 30 minutes, after which their color was compared against a negative control (yellow) and positive control (dark green) containing a mosquito which was fed with a 20% fructose solution.

#### Quantitative carbohydrate assays

Following this, glass tubes containing the cold anthrone samples were filled with anthrone reagent to a 5 mL mark, heated for 17 min at 92°C in a dry bath, and then cooled before being vortexed for 15-20 sec. A sample without a mosquito was prepared additionally as a control and blank for the spectrophotometer (Perkin Elmer Lambda 20 UV/Visible Spectrophotometer). The optical density (OD) of each sample was then determined at 625 nm. For samples with an OD_625_ above one, 200 μL of sample was diluted with 800 μL anthrone reagent that had been heated as above, giving a dilution factor of 5. The carbohydrate content was quantified using the OD values and a calibration line that had been created by performing the above procedure with samples containing 25, 50, 100, 150, and 200 μg of glucose solution. An ANOVA was used to compare the carbohydrate contents between the different groups using the software R [30].

### 2.5. Plant DNA Barcoding Assay

#### Mosquito trapping

Throughout the field season of 2021 (Late May – End of October), mosquitoes were collected weekly from ten residential yards in Blacksburg, VA (USA) containing a high diversity of ornamental and wild plant species. Mosquitoes were captured once a week using BG-2 Sentinel Traps (BioQuip) baited with an attractive lure and carbon dioxide released by dry ice placed in an adjacent cooler (BioQuip). The traps were deployed late afternoon and retrieved mid-morning (total: 16 - 17 hrs), which prevented mosquitoes from drying out or digesting crop sugars, which usually occurs within 24 hours after ingestion [26]. To further limit sugar digestion and degradation, mosquitoes were transported from the field to the lab on ice and stored at −70°C.

#### Mosquito sample processing and DNA extraction

Mosquitoes collected from the field were identified by morphological traits under a microscope [31], and washed in phosphate-buffered saline (Sigma-Aldrich, #806552-1L) to remove plant material contaminants, after which they were crushed in a microcentrifuge tube containing 100 μL of 0.3 M sodium acetate (Thermo Scientific, AM9740) and 200 μL of absolute molecular grade ethanol (Sigma-Aldrich, E7023-500mL) as in Wanjiku *et al.* (2021) [32]. After incubation for 30 minutes at −20°C, homogenates were centrifuged at 4°C for 10 minutes at 12,000g. Following centrifugation, 200 µL of the sample supernatant was tested for the presence of fructose using the cold anthrone method described above. After a 24-hour drying period, pellets were extracted using the RED Extract-N-AMP plant DNA extraction kit (Sigma-Aldrich, XNAP-1KT), according to the manufacturer’s instructions. Leaves from goldenrod collected in the field had their DNA extracted in a similar fashion and were used as a positive control.

#### PCR and Sanger sequencing

DNA was extracted from samples that gave a positive result for the anthrone test and genes of interest were amplified using PCR. Each sample was composed of 200 ng of DNA, PCR grade water (Fisher, AM9935), 10 μL of MyTaqHSmix (Bioline BIO-25045), 0.4 μM of forward and reverse primers targeting the chloroplast ribulose-1,5 biphosphate carboxylase/oxygenase large chain gene (*rbcLa*) (Primer R: GCTTCGGCACAAAAKARGAARCGGTCTC; Primer F: TATGTAGCTTAYCCMTTAGACCTTTTTGAAGA) or *trnh-psbA* intergenic spacer region (*trnH*) (*trnH*: CGCGCATGGTGGATTCACAATCC; *psbA*: GTTATGCATGAACGTAATGCT) to reach a sample volume of 22 μL. PCR cycling conditions for the *rbcLa* barcode were conducted as in Wanjiku *et al.*, 2021 with an initial denaturation of 95 °C for 1 min, 35 cycles of denaturation at 95 °C for 15 s, annealing at 50 °C for 40 s, and extension at 72 °C for 1 min followed by one cycle of final extension at 72 °C for 10 min. For the *trnH* barcode, cycling conditions followed Nyasembe *et al.*, 2018 [33] with a denaturation of 94 °C for 1 min, followed by 45 cycles of denaturation at 94 °C for 1 min, annealing at 55 °C for 40 s and 72 °C for 1 min, and a final extension at 72 °C for 10 min. 1% agarose gels and a 1 Kb DNA ladder (Genesee Scientific, 42-432) were used to visualize and confirm the size of the PCR products. Samples with clear, single bands of the correct size on the agarose gel were sent to Genewiz (South Plainfield, NJ, USA) for purification and Sanger sequencing. Sequences were then assembled using CLC main workbench (Qiagen) and compared to the GenBank database using the Basic Local Alignment Search Tool (BLAST) to search for candidate host plant species; only sequencing results with more than 85% homology were considered as potential candidates.

## 3. Results

### 3.1 Plant Visitation Assays

#### Landings

*Aedes aegypti* exhibited a high number of visitations for the goldenrod and yarrow ornamental plant species (Fig. 1B). Goldenrod had the highest number of landings from *Ae. aegypti*, while red impatiens and marigold had the least (one male “M” and one female “F”, respectively). We did not notice differences in landing activity between *Ae. aegypti* males and females, with the exceptions being butterfly bush (68.7% M, 31.3% F), Mexican heather (77.7% M, 22.3% F), and goldenrod (43.3% M, 56.7% F). Across all visitation assays, *Ae. aegypti* landed less on the plants compared to *Ae. albopictus*. Additionally, we found a greater variation in landing activity between *Ae. albopictus* males and females, compared to *Ae. aegypti*. Female *Ae. albopictus* landed more on the wave petunia (36.7% M, 63.3% F), marigold (33.3% F, 66.7% M), *Lantana* (12.4% M, 87.6% F), and *Scaevola* (22.3% M, 77.7% F). Conversely, males landed more on the *Celosia* (79.1% M, 20.9% F), butterfly bush (83.5% M, 16.5% F), goldenrod (78.6% M, 21.4% F), and yarrow (78.6% M, 21.4% F). Only one male and one female *Ae. albopictus* landed on *Guara* and Mexican heather. These two plant species also had the least total landings from *Ae. albopictus*, while wave petunia had the most.

#### Feedings

We observed *Ae. aegypti* feeding on five ornamental plant species: butterfly bush, *Guara*, *Lantana*, goldenrod, and yarrow (Fig. 1C). Female *Ae. aegypti* fed more than males; only females fed on butterfly bush (0% M, 100% F), *Guara* (0% M, 100% F) and *Lantana* (0% M, 100% F), and goldenrod (22.3% M, 77.7% F) had significantly more feedings from females. We did not observe *Ae. aegypti* feeding on the wave petunia, red impatiens, marigold, *Celosia*, or *Scaevola* which coincided with the low number of landings observed (Fig. 1B). Overall, *Ae. albopictus* fed more often than *Ae. aegypti*, which correlates with the higher number of landings from this species. In contrast to *Ae. aegypti*, we observed more total feeding events from male *Ae. albopictus* compared to females. *Celosia* (100% M, 0% F), butterfly bush (80.1% M, 19.9% F), goldenrod (83.5% M, 16.5% F), and yarrow (100% M, 0% F) all had a greater number of males’ feeding. Females fed more on the remaining four plants: marigold (33.5% M, 66.5% F), *Guara* (0% M, 100% F), *Lantana* (24.8% M, 75.2% F), and *Scaevola* (0% M, 100% F). We did not observe feeding from either mosquito species on Mexican heather, which corresponded with the low landing activity noted (Fig. 1B). Across all visitation assays, *Ae. aegypti* and *Ae. albopictus* fed the most often on goldenrod.

#### Survival

Across nine of the eleven ornamental species, *Ae. aegypti* females exhibited a higher average survival rate than males (Fig. S1A). For the remaining two ornamental species *Celosia* and *Lantana*, *Ae. aegypti* males and females shared an identical average survival rate at 97% and 80% survival, respectively. The lowest survival rates from both sexes were observed with the yarrow plant, with a 47% and 63% survival rate from males and females, respectively. For all ornamental species tested, *Ae. albopictus* females survived at a higher rate than males (Fig. S1B). The difference in survival rate between *Ae. albopictus* females and males was generally greater than *Ae. aegypti*. Survival rates for *Ae. albopictus* females and males were highest for the visitation assays with *Celosia* (97%) and *Scaevola* (80%), respectively; in contrast, the lowest rates occurred with marigold (63%) and *Lantana* (30%) for females and males, respectively.

### 3.2 Carbohydrate content assays

#### Qualitative analysis

Mosquitoes that tested positive for fructose using the cold anthrone test confirmed the presence of sugar feeding activity (Fig. 1E). Overall, *Ae. albopictus* exhibited a higher proportion of sugar feeding compared to *Ae. aegypti* (6.5% and 3.3%, respectively). Differences in sugar feeding proportions were observed between mosquito species for most ornamental plant species, with *Ae. albopictus* testing positive only with *Scaevola*, goldenrod, and yarrow, and *Ae. aegypti* testing positive only with butterfly bush and *Lantana*. No positive tests from either species were found for marigold and Mexican heather. We observed the highest sugar feeding rate from *Ae. albopictus* and goldenrod (20.9%), followed by *Ae. albopictus* with *Celosia* (18.2%) and *Ae. aegypti* with *Lantana* (12.5%). For some assays, such as goldenrod with *Ae. aegypti*, multiple feeding events were observed but no mosquitoes tested positive for fructose consumption.

#### Quantitative analysis

The total amount of carbohydrates among the fructose-positive mosquitoes was quantified using the warm anthrone protocol (Fig. 1F). We found the highest carbohydrate concentration values in a female *Ae. albopictus* (541.43 µg ± 124.90 µg) and a male *Ae. aegypti* (301.43 µg ± 100.43 µg) after the visitation assay with the wave petunia, followed by a female *Ae. aegypti* with *Lantana* (249.29 µg ± 39.71 µg). Generally, higher carbohydrate concentrations were recorded from *Ae. aegypti* individuals compared to *Ae. albopictus*. The total carbohydrate concentrations between mosquitoes testing positive for fructose consumption and mosquitoes testing negative were also compared (Fig. S2A and S2B). For both mosquito species, concentrations from negative-testing mosquitoes were consistently lower than positive-testing individuals. However, due to the low number of samples for “positive” mosquito groups, we could not demonstrate the higher average contents in carbohydrates in the “positive” groups with statistical significance (for all comparisons, Student *t*-tests, p > 0.05).

### 3.3 GC-MS Analysis of Plant Odor

Headspace collections and GC-MS analyses were then conducted to determine the VOC composition of the scent of each ornamental species (Fig. 2, Table 1). We found that terpenoids including linalool and aliphatics such as nonanal were in abundance across each ornamental species tested. Nonanal had the highest relative concentration in the wave petunia (44.1%), red impatiens (35.7%) and *Guara* (57.8%). Other aliphatic compounds, including 2-hexanal and santolina triene, were present in small concentrations in all ornamental species except goldenrod. In *Lantana*, Mexican heather and *Scaevola*, *ß*-ocimene comprised the majority of the scent profile, representing 42.7%, 79.2%, and 56.9% of the scent composition, respectively. Benzaldehyde and limonene were found frequently across most of the ornamental species at varying concentrations. Benzaldehyde was found at its highest relative abundance in *Lantana* (29.6%), while limonene was found at high concentrations in the scent of marigold (17.7%) and goldenrod (21.6%). Some less common terpenoid volatiles were unique to certain ornamental species, yet still displayed a major relative peak abundance in their associated plant scent. For example, *ß*-bisabolene was present only in the scent of *Celosia* but exhibited the highest relative peak abundance (30.8%). Germacrene D was a major constituent of the goldenrod scent (15%), and was present at smaller concentrations in marigold, *Lantana* and yarrow. We found that *ɑ*-pinene was in high abundance in the scent of goldenrod (25%) and was one of most abundant compounds in marigold (16.3%) and yarrow (7.7%). *ɑ*-farnesene was abundant only in the butterfly bush, making up 87.8% of the total scent’s peak area abundance. *ß*-phellandrene was found at small concentrations in goldenrod, *Lantana* and marigold, but had the highest relative peak abundance in yarrow (28%), with cis-verbenol being the second most abundant compound (26.8%). Finally, caryophyllene was the dominant compound in the scent profile of the marigold (20.5%) and was present at smaller concentrations in *Scaevola* (8.4%) and yarrow (3.2%).

**Fig. 2.**
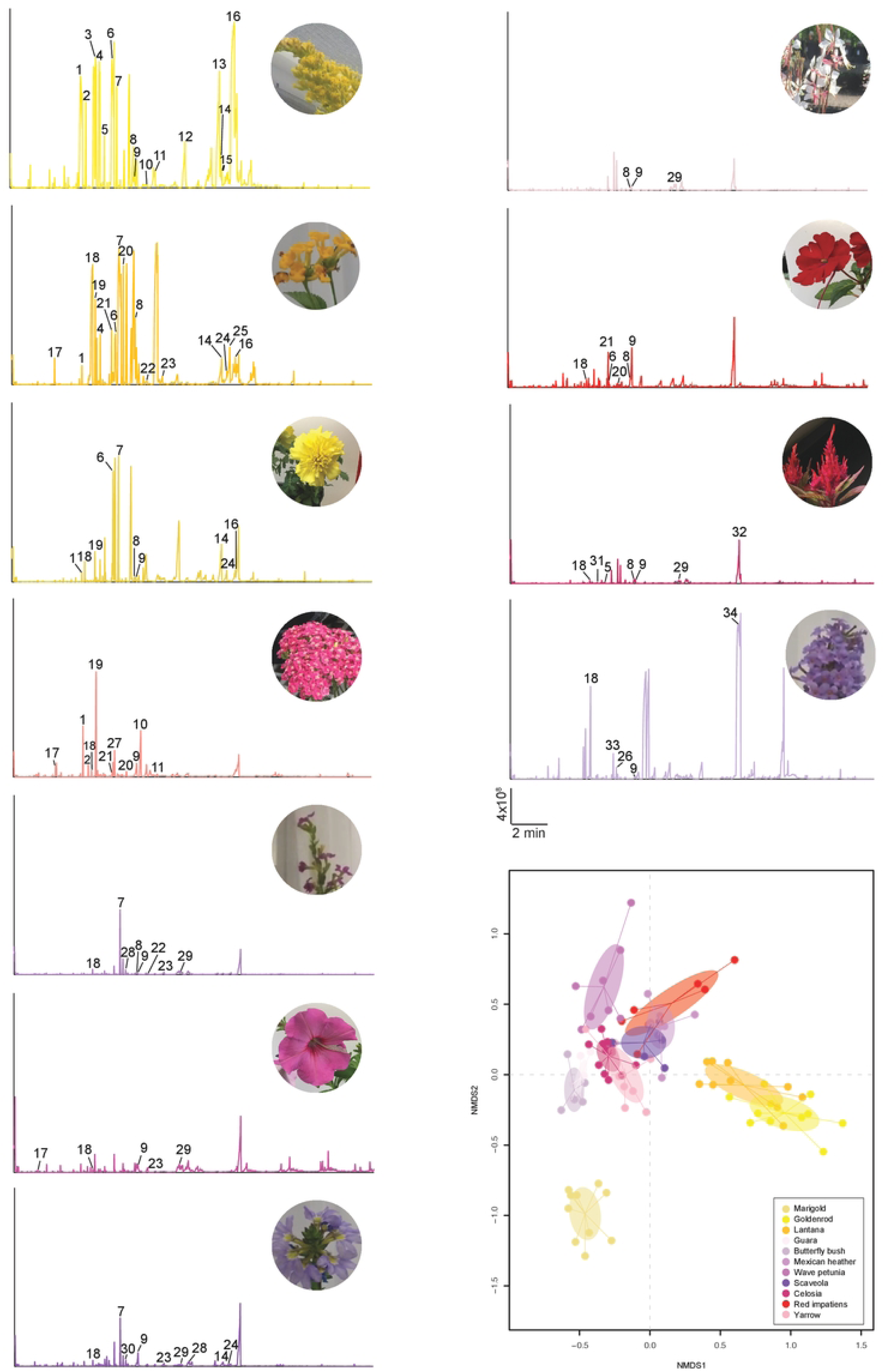
GCMS analyses of the floral volatiles emitted by eleven ornamentals. Pictures of the ornamental species are displayed on the right of each chromatogram. Numbers in the chromatograms correspond to: 1, *ɑ*-pinene; 2, camphene; 3, *β*-pinene; 4, *β*-myrcene; 5, *ɑ*-phellandrene; 6, D-limonene; 7, *β*-ocimene; 8, linalool; 9, nonanal; 10, verbenol; 11, endo-borneol; 12, bornyl acetate; 13, aristolene; 14, caryophyllene; 15, γ-elemene; 16, germacrene D; 17, hexenal; 18, benzaldehyde; 19, *β*-phellandrene; 20, γ-terpinene; 21, p-cymene; 22, myroxide; 23, methyl salicylate; 24, *β-*farnesene; 25, humulene; 26, benzeneacetaldehyde; 27, eucalyptol; 28, m-ethylacetophenone; 29, p-cymen-7-ol; 30, γ-chlorobutyrophenone; 31, 5-octen-1-ol (Z); 32, *β-*bisabolene; 33, benzyl alcohol; 34, *ɑ*-farnesene.

We conducted a NMDS (non-metric multidimensional scaling) analysis to compare the scent profiles of each ornamental plant species based on the chemical compounds and their relative abundance (Fig. 2). Most of the ornamental species exhibited a distinct clustering of samples, indicating a unique and distinct scent composition (ANOSIM, R = 0.93, *p* = 0.001) (stress = 0.1). Of note, marigold, *Lantana* and goldenrod samples showed strong individual clustering; according to our GC-MS analysis, the scent profile of these three plants contains volatile compounds that are highly abundant relative to other ornamentals. Interestingly, goldenrod and *Lantana* samples were clustered close together, while the marigold samples were isolated in the bottom-left quadrant of the NMDS. This positioning also reflects the behavior of the mosquitoes as neither species obtained carbohydrates from marigold, thus supporting the hypothesis that mosquitoes respond to ornamental species based on their scent profile.

### 3.4 Plant DNA Barcoding

#### Carbohydrate analysis

A total of 2360 mosquitoes were collected during the field collection season. We identified a total of five species: *Ae. albopictus* (N = 1527; 54.5%), *Culex pipiens* (N = 574; 24.3%), *Anopheles punctipennis* (N = 29; 1.2%), *Ae. vexans* (N = 93; 3.9%), and *Ae. triseriatus* (N = 96; 4.1%). The number of mosquitoes collected varied by week, with a large peak occurring in mid-July that gradually decreased until the end of the season in late October (Fig. 3A). The overall fructose positivity rate was 38% (N = 903 positive tests; Fig. 3B). Throughout the season, this rate generally ranged between 25% and 50% but did not gradually decrease, even towards the end of the season in October (Fig. 3C). In addition, the positivity rate varied across trap sites and ranged between 26% and 46% (Fig. 3D). The majority of mosquitoes captured were females (females: N = 1992; 84%; males: N = 368; 16%). However, we found that males had a higher proportion of positive sugar tests (N = 164; 45%) compared to females (N = 739; 37%) (Fig. 3B). The total number of *Ae. albopictus* mosquitoes captured across trap sites ranged between 40 and 161, with the exception of trap site 8, where the highest number of *Ae. albopictus* mosquitoes were trapped (N = 670; Fig. S3C). The proportion of *Ae. albopictus* testing positive for fructose consumption ranged between 24% and 50% across the ten trap sites (Fig. S3B).

**Fig. 3.**
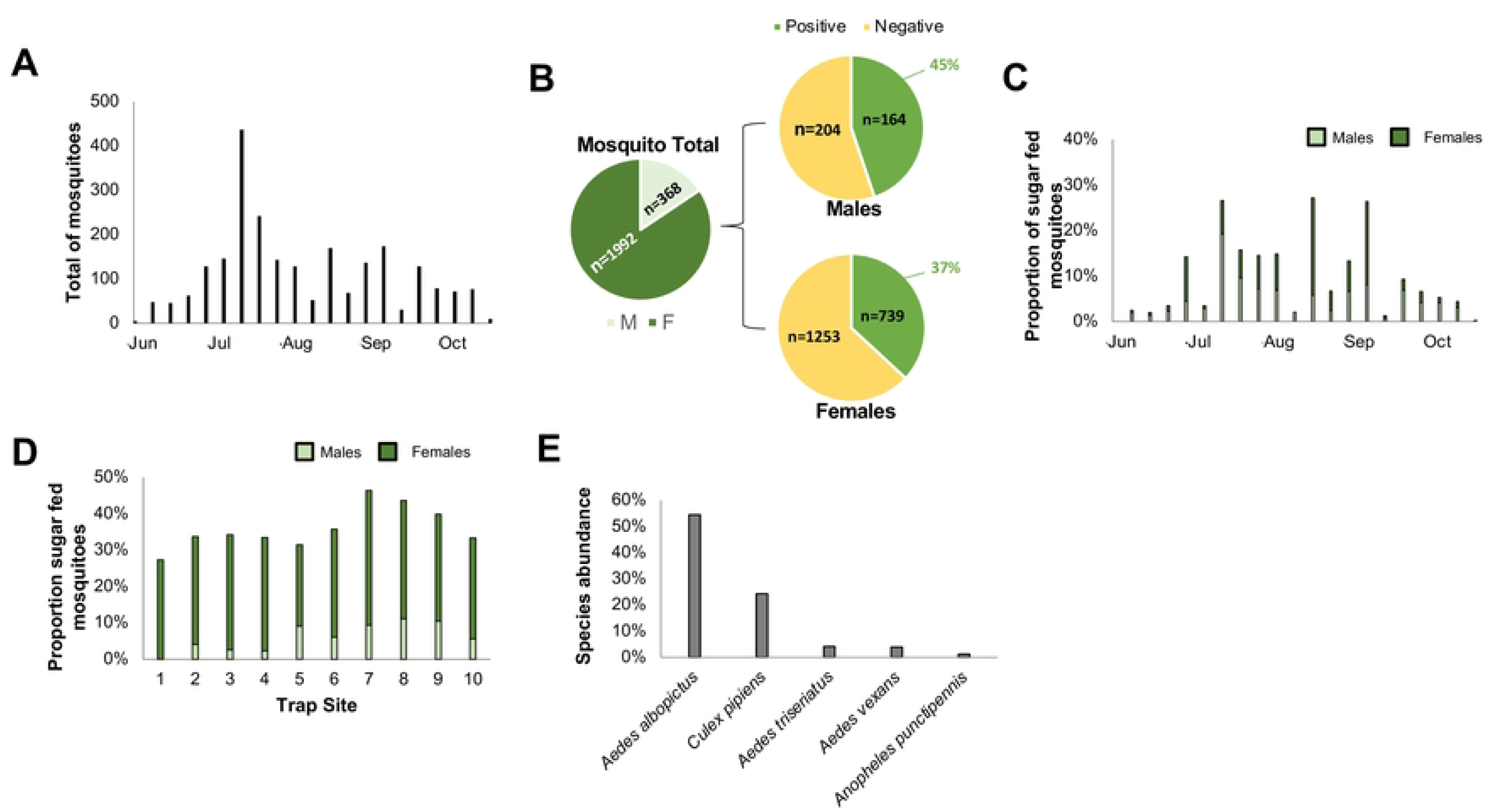
Mosquito sugar-feeding activity in residential Blacksburg, Va. (A) Total mosquitoes captured across the ten trapping locations per week of the candidate plant host study. (B) The proportion of males (light gray) and females (dark gray) captured across the trapping locations (*left*) and the proportion of positive sugar-feeding tests for all males and females. (C) The average proportion of positive sugar-feeding tests per week of the field season for males (light green) and females (dark green). (D) The average proportion of positive sugar-feeding tests across the field season for each trap site. (E) The proportion of different mosquito species captured across all trapping locations.

#### Candidate host plants

A total of 75 mosquitoes contained plant DNA that was amplified using the *rbcLa* primers while 144 samples were amplified using *trnH* primers. From these 219 amplifications, we identified 26 unique candidate host plants from 18 different families (Table 2). Interestingly, most of these plants are commonly planted as ornamentals in gardens (*e.g.*, *Prunus spp., Acer spp., Carya illinoinensis*). Eighteen of the 26 candidate host plant seqeuences were fully assembled; the remaining 8 did not assemble and were identified based on the forward or reverse primer sequences. Most of the plant candidates were identified from DNA that was extracted from *Ae. albopictus* females. However, some were also determined for male *Ae. albopictus* and female *Cx. pipiens*, *An. punctipennis*, *Ae. japonicus* and *Ae. vexans*. Plant species that were identified from the crops of multiple mosquitoes include *Trifolium repens* (*rbcLa, trnH*; N = 12), *Prunus spp.* (*rbcLa*; N = 18)*, Drosera filiformis* (*rbcLa*; N = 2)*, Plantago spp.* (*trnH*; N = 5) and *Acer spp.* (*rbcLa*, *trnH*; N = 4). Most plant species were identified from mosquitoes during specific weeks of the season (Table 2). For example, we found that mosquitoes fed on *Oxalis dillenii* in July and on *Drosera filiformis* only in August. However, species like *Trifolium repens* and *Prunus virginiana* were identified from mosquitoes captured throughout the entire field season.

## 4. Discussion and Perspectives

The first part of this work sheds light on the interaction between *Ae. aegypti* and *Ae. albopictus* mosquitoes and ornamental plants and provides some insights into the understudied field of mosquito phytophagy. Through behavioral observations coupled with crop sugar screening, we found a clear preference for specific ornamental plants over others which was unique to each mosquito species. In addition, GC-MS analyses allowed us to identify the VOCs present in each flower’s scent, building on our current understanding of critical VOCs that mosquitoes may use to locate their sugar meals. We also examined the potential effect that flower color, morphology and nectar accessibility might have on mosquito sugar feeding.

For both mosquito species, visitation and sugar feeding activities varied significantly between each ornamental flower. High numbers of landings and feedings were seen for both mosquito species on the goldenrod. Occurrences of mosquitoes visiting goldenrod have been recorded as early as 1907 [34], and multiple genera have been observed visiting the flower in the field (personal observations). One stalk of goldenrod can contain hundreds of shallow flowers filled with nectar, presumably making it easy for mosquitoes to access the flowers with their proboscis (*i.e* piercing mouthpart) and feed. Interestingly, close to 25% of *Ae. albopictus* tested positive for fructose consumption in the present study, yet no *Ae. aegypti* gave positive tests, implying that feeding on goldenrod may be species-specific. *ɑ*-pinene, the most abundant compound in goldenrod’s scent, has been tested against *Ae. aegypti* and *Ae. albopictus* for larvicidal or mosquitocidal effects [35,36]. However, its role in mediating attraction or repellency in mosquitoes remains unclear. In addition, Germacrene D was present in high abundance (15.1%). This volatile has been shown to repel *Ae. albopictus* when tested in bioassays [37]. However, the ratio of odorants in a given scent is critical for plant-host location by mosquitoes [5,38]. The ratio of Germacrene D is likely important for mediating attraction in the diverse VOC blend of goldenrod’s scent, and this may be the case for other VOCs that are repellent on their own but found in potentially attractive plants throughout this study.

Interestingly, we observed high visitation activity from both mosquito species on yarrow, which has generally been defined as a mosquito-repellent flower [39]. Although a high number of feedings was observed from *Ae. albopictus*, low nectar consumption rates were found in the total carbohydrate assays. Moreover, *Ae. aegypti* did not obtain any carbohydrates from their observed feeding. Though it seems to provide a low nutritional reward, it is clear that this flower is not aversive to mosquitoes, as shown by the frequent landings we observed from both species. The scent of this plant was dominated by cis-Verbenol and *ß*-phellandrene. When mixed with (-)-limonene, *ß*-phellandrene was found to be repellent against *Ae. albopictus* mosquitoes [37]. It was also shown that cis-verbenol is present in essential oils that exhibit larvicidal activity against *Ae. albopictus* and *Ae. aegypti* [40]. However, the capacity of these compounds to attract or repel *Ae. aegypti* and *Ae. albopictus* has yet to be tested individually. In addition to yarrow, we observed high visitation activity from *Ae. albopictus* on *Celosia*, marigold, and wave petunia. More *Ae. albopictus* landed and fed on *Celosia* compared to *Ae. aegypti*. This may be in part due to *β*-bisabolene dominating the scent of this plant, which was found to be present in essential oils that repel *Ae. aegypti* [41]. Almost 20% of *Ae. albopictus* in plant visitation assays with *Celosia* tested positive for fructose consumption, but contained a low concentration of total carbohydrates, possibly indicating low nectar content in the flowers. Similarly, we observed high landing and feeding activity from *Ae. albopictus* on marigold, but no mosquitoes tested positive for fructose consumption. This could indicate low nectar content or can be attributed to the morphology of *Celosia* (tall, feathery plumes) and marigold (wide, bowl-like) flowers not being adapted for a mosquito’s proboscis to probe and feed on. Interestingly, caryophyllene was the most abundant compound of marigold’s scent and is a major constituent in an essential oil repellent to *Ae. aegypti* but has not been tested with *Ae. albopictus* [42]. Mosquito response to caryophyllene is likely species-specific, as we observed multiple landings and feedings from *Ae. albopictus* and only one landing from *Ae. aegypti*.

We did not observe feeding activity by either species on wave petunia, yet individuals of both species tested positive for nectar consumption and exhibited the highest carbohydrate concentrations in this study. It is, however, possible that mosquitoes fed on stems / leaves containing a high concentration of sugars either outside of the camera’s frame, or outside of the camera’s recording time at night when petunias emit a stronger scent [1,43,44]. We identified *β*-ocimene as the most abundant volatile in the *Lantana* headspace, which has been shown to elicit dose-dependent repellent responses in multiple aedine species, including *Ae. aegypti* [33,45]. Yet, multiple landings were observed from both species in the present study. This could be attributed to the ratio of different volatiles in *Lantana’s* scent profile; benzaldehyde was also a major constituent in *Lantana* scent and has been hypothesized to attract mosquitoes when present in volatile blends [46]. It has been shown that *Ae. albopictus* oviposit in water buckets near flowering butterfly bushes, but this may not be indicative of a role as a sugar source due to the lack of positive fructose tests seen for *Ae. albopictus* here [47]. *ɑ*-farnesene was a prominent volatile in the scent profile of butterfly bush; its enantiomer β-farnesene has demonstrated a low degree of repellence in *Ae. aegypti* mosquitoes [48]. However, mosquitoes have been shown to have enantioselectivity for odor compounds, so *β*-farnesene may not be representative of a response to *α*-farnesene [49] as implied by the high number of attempted feedings observed from both mosquito species.

The ornamental plants with the lowest visitation activity were *Guara*, Mexican heather and *Scaevola*. As with wave petunia, we observed a low number of feedings from both mosquito species on *Guara*, yet several mosquitoes tested positive for sugar feeding. This may be due to high concentrations of nonanal, a VOC known to play a role in attracting *Ae. aegypti* to orchids, emanating from both plants at a time outside of the camera recording [7]. *β*-ocimene is a compound with a high degree of ubiquity in plant scents (observed in 75% of 63 plant families) and was the most abundant compound in the scents of Mexican heather and *Scaevola* [50]. As mentioned above, this VOC is repellent to *Ae. aegypti* at certain concentrations and may be deterring mosquitoes from landing on these flowers. Some *Ae. albopictus* were observed feeding on *Scaevola* and tested positive for sugar consumption, but no mosquito was seen feeding on nor tested positive for crop nectar with Mexican heather. This may be due to the large proportion of *β*-ocimene dominating the odor profile of Mexican heather (79.24%) relative to other ornamentals (56.93% in *Scaevola* and 42.72% in *Lantana*).

After determining that a preference for ornamental flowers exists in mosquitoes, we expanded our focus from the laboratory to the field where ornamental plants are in high abundance to determine which ornamental species local mosquitoes feed on. Using plant DNA barcoding, we identified 26 unique plant species that various mosquito species feed on in an urban environment and noticed variations depending on the time of year. When using *rbcLa* primers, the genus that appeared most often was *Prunus*, a group of trees and shrubs that are cultivated for their fruits and decorative qualities and was identified from the crop of both male and female *Ae. albopictus* and female *Cx. pipiens*. For each *Prunus* result, several species consistently appeared with the same high percent identity coverage, indicating that the *rbcLa* gene is highly conserved amongst *Prunus* species. *Prunus* species identified from BLAST present in the study area include *Prunus virginiana* (“bitterberry” or “chokecherry”) and *Prunus padus* (“bird cherry”); *Prunus virginiana* is native to North America while *Prunus padus* is planted as an ornamental and is native to northern Europe and Asia. It is possible mosquitoes are obtaining nectar from the plants’ flowers, but it is possible they feed on the small berry-like fruits produced by both plants.

*Trifolium repens* was identified frequently throughout the season using both the *rbcLa* and *trnH* barcodes and was targeted almost exclusively by female *Ae. albopictus* mosquitoes. This plant is extremely common in lawns and along roadsides in the United States and produces a substantial amount of nectar that gathers at the bottom of tube-shaped florets that are assumedly shallow enough for a mosquito’s proboscis to reach [51,52]. It was identified from some plant DNA samples using both barcodes, providing robust evidence that local invasive mosquitoes were using it as a sugar source. In a recent study by Kicel *et al.* (2010), 1-octen-3-ol was determined as a major constituent in the scent profile of *Trifolium repens’* stems and leaves [53]. This compound also emanates from human skin and has been reported as a kairomone (*i.e.*, attractant) for *Aedes* mosquito species (*Ae. albopictus* included), especially when acting synergistically with CO_2_ [54,55]. Due to the prevalence of *Trifolium repens* in urban locations containing an abundance of hosts, *Aedes* mosquitoes are likely responding to both 1-octen-3-ol and CO_2_ when host-seeking in these areas [56]. In addition, occurrences of *Trifolium repens* have been documented in every continent, and it is able to grow in a range of climates from arctic to tropical, potentially contributing to *Ae. albopictus*’ wide establishment as an invasive disease vector [57,58].

Another plant host that appeared several times in our results using both barcodes was the maple tree (*Acer spp.*). Maple trees are some of the most frequently planted ornamental trees in North America [59]. These trees do produce flowers and generally bloom from March to June. Interestingly, mosquitoes containing maple tree DNA were captured in July and August. Therefore, it is possible that the captured *Ae. albopictus* mosquitoes obtained carbohydrates by feeding on extra floral nectaries or from the leaves of the maple [1,44]. Alternatively, mosquitoes have been observed to feed on phloem sap, and may be doing so in this case due to the maple tree’s high sap production [60, 61]. An important point to consider is that both *Ae. albopictus* and *Acer spp.* are native to eastern Asia; it is thus possible that *Ae. albopictus* has been feeding on maple trees in its native habitat. Due to its commonality in urban settings, maple trees may also be contributing to this mosquito’s success in establishing itself as an invasive species.

Mosquitoes have been reported feeding on plant fruits in as early as 1758 [62]. Observations of mosquitoes feeding on pears and watermelons have been made in the 1900s, and the data presented here supports this with observed amplifications of *Pyrus communis* (pear) and *Citrullus mucosospermus* (watermelon) DNA [63]. To add to these observations of fruit-bearing plants, we identified *Solanum lycopersicum* (tomato), *Cucumis sativus* (cucumber), and *Cucurbita pepo* (pumpkin) as mosquito plant hosts. To our knowledge, this is the first published evidence that *Ae. albopictus*, or any mosquito species may feed on these plants. Furthermore, these plant hosts were identified only from mosquitoes captured in August and September, highlighting the importance of phenology studies for establishing mosquito sugar feeding activity in the wild. Previous studies have shown *An. arabiensis* larvae will feed on the pollen of maize plants in sub-Saharan Africa; gravid female *An. arabiensis* are stimulated by volatiles released by maize pollen and lay their eggs near it so as to provide food for their offspring [64]. In the same study, an attractant was successfully developed using a blend of volatiles from the scent of maize pollen. In the present study, *Cx. pipiens* used *Zea mays* (maize) as a plant host, indicating that the attractive blend could potentially be used as a multi-species attractant.

This study also provides a comparison for the different plant families that *rbcLa* and *trnH* barcodes can identify. Success rates in species discrimination have been shown to vary between these barcodes, and the present study shows variations in the plant families they can identify [65,66]. When comparing the ability of *rbcLa*, *trnH*, and *matK* to accurately identify legume (Fabaceae) species, *trnH* ranked the highest and was only slightly more accurate than *rbcLa* [67]. In the present study, *trnH* identified a higher amount of legume species, but had a similar percent identity to legume DNA sequences amplified by *rbcLa*, supporting the conclusion from Sanchez *et al.* (2020). A similar study was done previously to compare barcodes in discriminating Cucurbitaceae species, and *trnH* was once again found to exhibit the highest accuracy [68]. The *trnH* barcode identified multiple Cucurbitaceae species in the present study, whereas *rbcLa* identified none. The *rbcLa* barcode identified 18 different DNA sequences belonging to the Rosaceae family and *Prunus* genus but could not discriminate these sequences at the species level. When comparing the performance of *rbcLa*, *matK*, rpoC1 and ITS2 barcodes in identifying Rosaceae species, a previous study’s results suggested that ITS2 is the most effective [69].

Nectar composition and availability are important factors to consider when analyzing the results obtained in the present study. Traditionally, plants modify the chemical composition of their nectar to serve as rewards for specific pollinators that they share a symbiotic relationship with [70]. Because mosquitoes are considered generalist pollinators, it is unlikely that any of these ornamentals are developing their nectar for mosquito attraction specifically [71]. The concentration and composition of compounds within the nectar of the eleven tested ornamentals and the plant host candidates determined by sequencing still play a part in how attractive the given plant is to the mosquito, but future work using liquid-chromatography mass-spectroscopy (LC-MS) may further help our understanding of the nutrients that mosquitoes obtain by visiting the plants.

Overall, this study highlights the importance of expanding our understanding of mosquito phytophagy, in particular for invasive species. We show that *Ae. aegypti* and *Ae. albopictus* can make use of energy provided to them from ornamental plant sugars which certainly help establish their presence in urban, heavily populated areas in the US. The data presented in this work can be shared with city officials and the general public to inform them of ornamental plants that may or may not be attracting mosquitoes to the public areas they are being planted. In areas with a higher risk of mosquito-borne disease transmission, the experiments described here can be used to enact field removal of plants fed on by mosquitoes and to develop baits / repellents to mitigate the spread of disease. Elucidating the mosquito-plant relationship is pivotal for maintaining the development of effective and environmentally friendly disease vector control techniques.

## Acknowledgements

We thank Darren Dougharty and Shajaesza Diggs for their support with mosquito colony care. We would also like to thank Danny Eanes for his technical support as well as the residents of Blacksburg who provided their backyards for mosquito collection. We are grateful to members of the Lahondère laboratory and Vinauger laboratory for their feedback on the present study. This research was funded by the Eppley Foundation for Research, the USDA NIFA (VA-160160 and 2020-68018-30674), the Department of Biochemistry, The Fralin Life Science Institute, the Global Change Center at Virginia Tech. The following reagent was obtained through BEI Resources, NIAID, NIH: *Aedes aegypti*, Strain ROCK, MRA-734, contributed by David W. Severson and Aedes albopictus, Strain ATM-NJ95, Eggs, NR-48979.

## Figure and table legends

**Table 1.** Concentrations of ornamental volatile compounds. Concentrations are represented as an average in ng / µL. The proportion of each compound relative to other compounds present in each scent is displayed as a percentage. *a,d-Gala-octonic phenylhydrazide.

**Table 2.** Candidate host plant sources as determined by barcoding and Sanger sequencing. The species of mosquito that the DNA was extracted from and the number of mosquitoes that contained that DNA is represented. The phenology of the candidates is displayed on the right; highlighted boxes indicate when the mosquito fed on the given candidate.

## Supplementary figure legends

**Figure S1.**
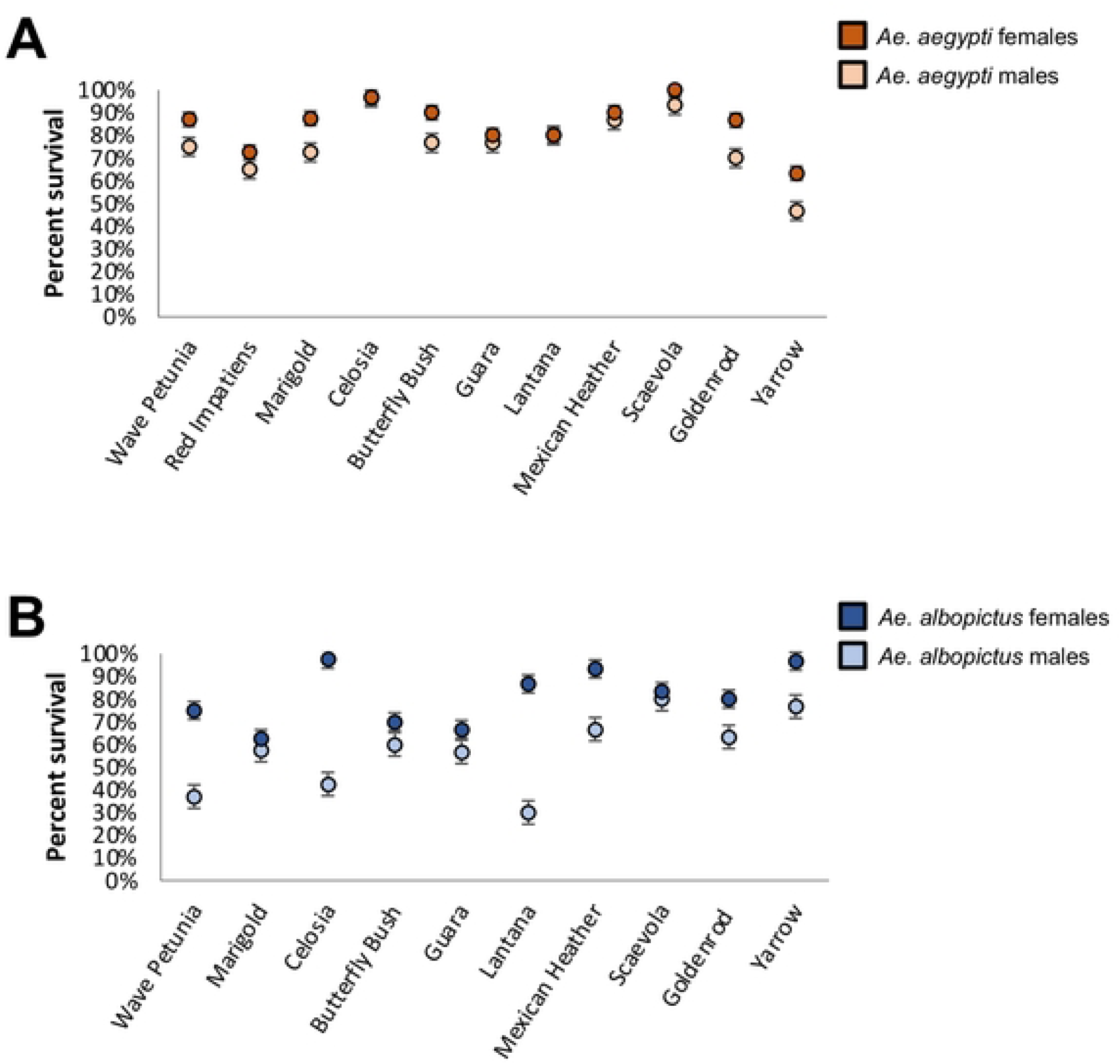
Survival rate of mosquitoes used for plant visitation assays (16 hrs; 9 a.m. - 5 p.m.). Percentages are represented as an average of the three replicates (n = 30 for each data point) performed for each plant species. (A) *Ae. aegypti* survival rate for males (light orange) and females (dark orange). (B) *Ae. albopictus* survival rate for males (light blue) and females (dark blue).

**Figure S2.**
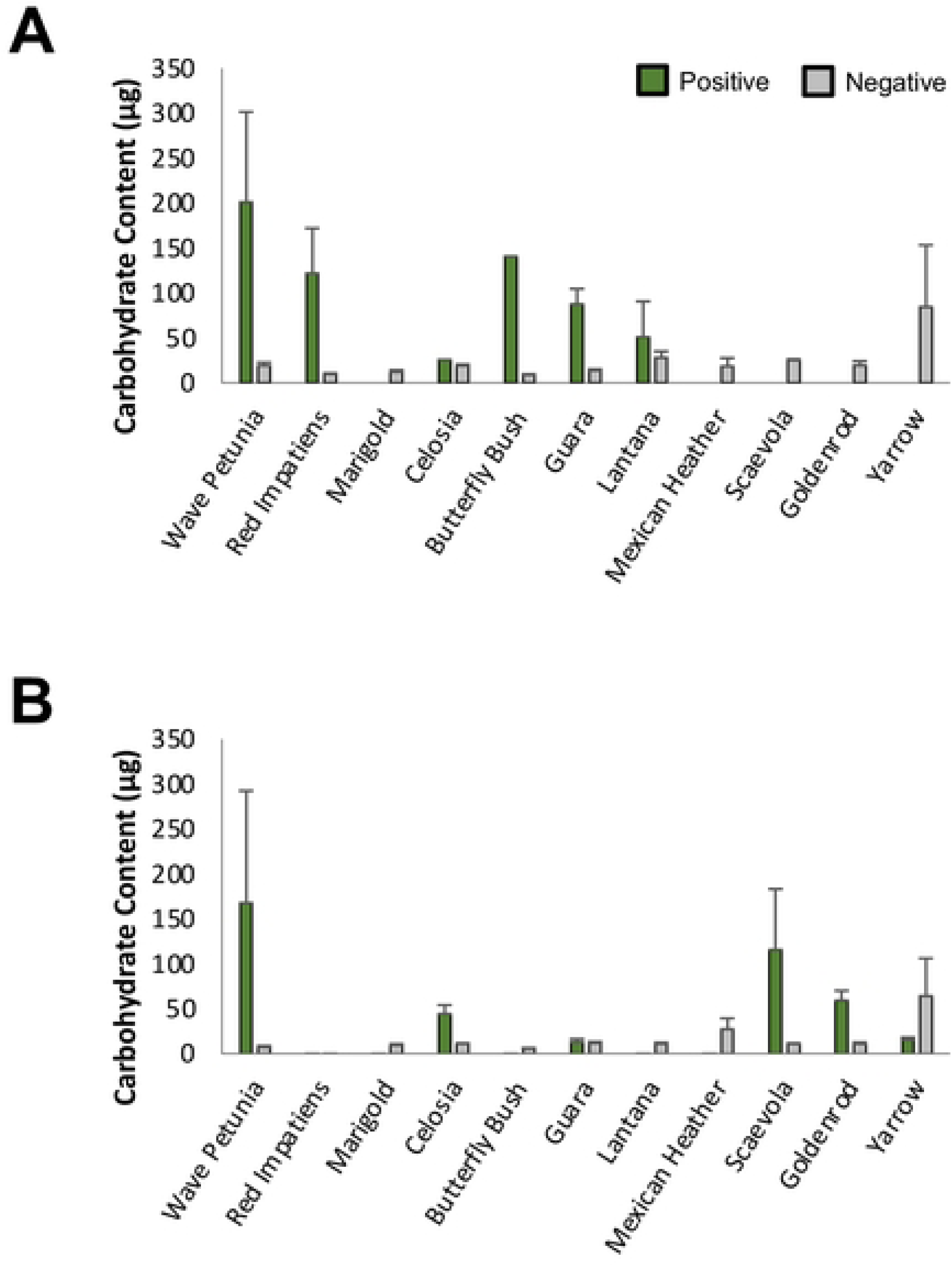
Total carbohydrate concentrations (µg) for (A) fructose-positive mosquitoes. Carbohydrate concentrations for fructose-positive and fructose-negative mosquitoes are also represented for (B) *Ae. aegypti* and (C) *Ae. albopictus*. Values are represented as averages of all mosquitoes that were alive following the plant visitation assays and could be measured for carbohydrates.

**Figure S3.**
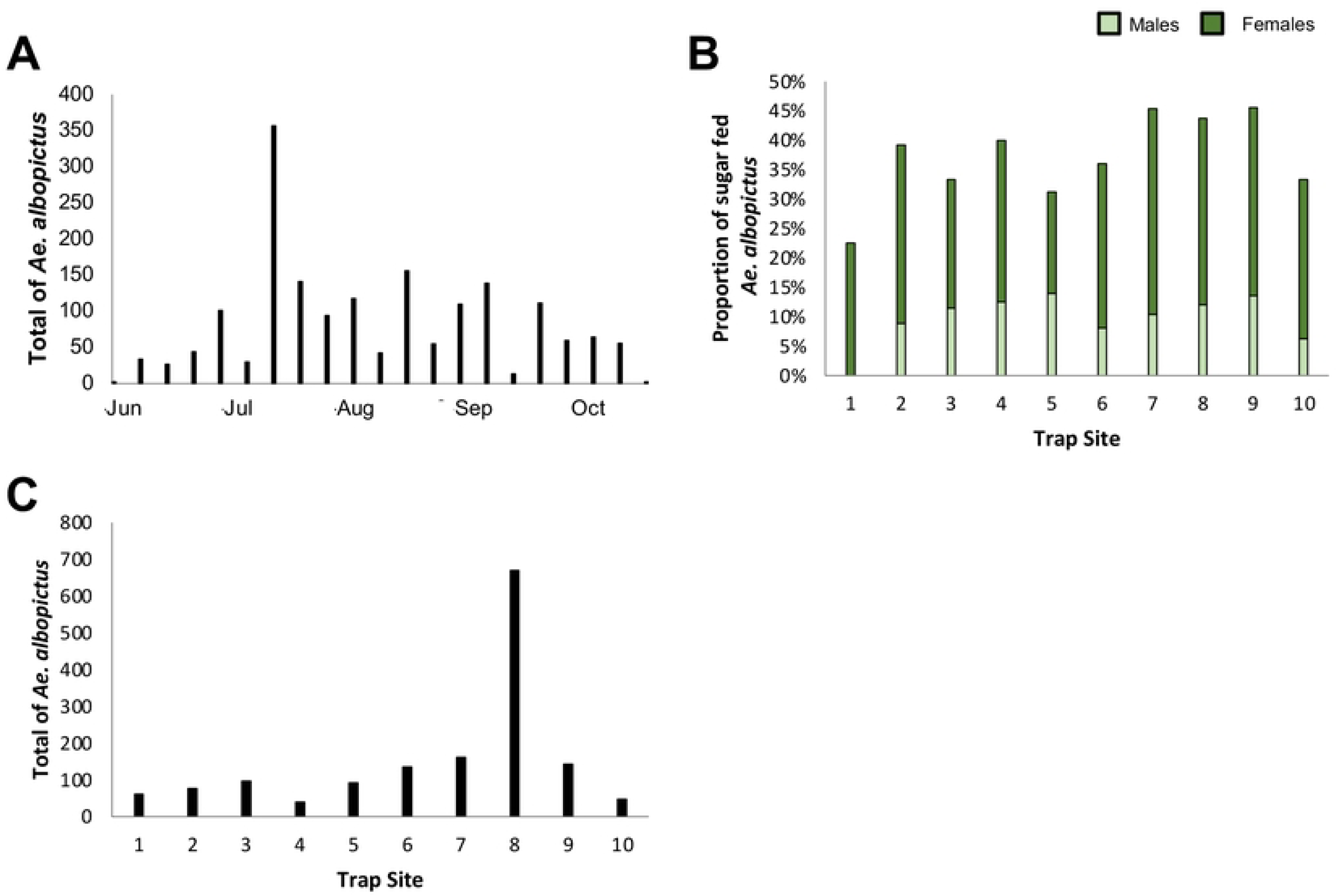
*Aedes albopictus* prevalence and sugar-feeding activity in residential Blacksburg. (A) The total number of *Ae. albopictus* captured per week during the candidate plant host study. (B) The proportion of positive sugar tests per trap site given by male (light green) and female (dark green) *Ae. albopictus*. (C) Total *Ae. albopictus* captured across each of the ten trapping sites.

## References

1. Foster WA. Mosquito sugar feeding and reproductive energetics. Annual Review of Entomology. 1995; 40(1): 443–474.

2. Baker H, Baker I. Amino-acids in Nectar and their Evolutionary Significance. Nature. 1973; 241: 543–545.

3. Nicolson SW, Thornburg RW. Nectar chemistry. In Nectaries and Nectar. 2007; 215–264. Springer, Dordrecht.

4. Rivera-Perez C, Clifton ME, Noriega FG. How micronutrients influence the physiology of mosquitoes. Current Opinion in Insect Science. 2017; 23: 112–117.

5. Nyasembe VO, Torto B. Volatile phytochemicals as mosquito semiochemicals. Phytochemistry Letters. 2014; 8: 196–201.

6. Nyasembe VO, Teal PE, Mukabana WR, Tumlinson JH, Torto B. Behavioural response of the malaria vector *Anopheles gambiae* to host plant volatiles and synthetic blends. Parasites & Vectors. 2012; 5: 234.

7. Lahondere C, Vinauger C, Okubo RP, Wolff GH, Chan JK, Akbari OS, Riffell JA. The olfactory basis of orchid pollination by mosquitoes. Proceedings of the National Academy of Sciences. 2020; 117(1): 708–716.

8. Jaleta KT, Hill SR, Birgersson G, Tekie H, Ignell R. Chicken volatiles repel host-seeking malaria mosquitoes. Malaria Journal. 2016; 15(1): 354.

9. Liu N. Insecticide Resistance in Mosquitoes: Impact, Mechanisms, and Research Directions. Annual Review of Entomology. 2015; 60: 537–559.

10. Gu ZY, He J, Teng XD, Lan CJ, Shen RX, Wang YT, Zhang N, Dong YD, Zhao TY, Li CX. Efficacy of orally toxic sugar baits against contact-insecticide resistant *Culex quinquefasciatus*. Acta Tropica. 2020; 202: 105256.

11. Khallaayoune K, Qualls WA, Revay EE, Allan SA, Arheart KL, Kravchenko VD, Xue RD, Schlein Y, Beier JC, Muller GC. Attractive toxic sugar baits: Control of mosquitoes with the low-risk active ingredient dinotefuran and potential impacts on nontarget organisms in Morrocco. Environmental Entomology. 2013; 42: 1040–1045.

12. Revay EE, Schlein Y, Tsabari O, Kravchenko V, Qualls W, De-Xue R, Beier JC, Traore SF, Doumbia S, Hausmann A, Müller GC. Formulation of attractive toxic sugar bait (ATSB) with safe EPA-exempt substance significantly diminishes the *Anopheles sergentii* population in a desert oasis. Acta Tropica. 2015; 150: 29–34.

13. Fiorenzano JM, Koehler PG, Xue RD. Attractive toxic sugar bait (ATSB) for control of mosquitoes and its impact on non-target organisms: A review. International Journal of Environmental Research and Public Health. 2017; 14(4): 398.

14. Bokore GE, Svenberg L, Tamre R, Onyango P, Bukhari T, Emmer Å, Fillinger U. Grass-like plants release general volatile cues attractive for gravid *Anopheles gambiae* sensu stricto mosquitoes. Parasites & Vectors. 2021; 14(1): 1–15.

15. Meza FC, Roberts JM, Sobhy I.S, Okumu FO, Tripet F, Bruce TJ. Behavioural and Electrophysiological Responses of Female *Anopheles gambiae* Mosquitoes to Volatiles from a Mango Bait. Journal of Chemical Ecology. 2020; 1–10.

16. Qualls WA, Müller GC, Traore SF, Traore MM, Arheart KL, Doumbia S, … Beier JC. Indoor use of attractive toxic sugar bait (ATSB) to effectively control malaria vectors in Mali, West Africa. Malaria Journal. 2015; 14(1): 1–8.

17. Fryzlewicz L, VanWinkle A, Lahondere C. Development of an Attractive Toxic Sugar Bait for the Control of *Aedes j. japonicus* (Diptera: Culicidae). Journal of Medical Entomology. 2022; 59(1): 308–313.

18. Müller GC, Junnila A, Schlein Y. Effective control of adult *Culex pipiens* by spraying an attractive toxic sugar bait solution in the vegetation near larval habitats. Journal of Medical Entomology. 2010; 47(1): 63–66.

19. Qualls WA, Xue R, Revay EE, Allan SA, Müller GC. Implications for operational control of adult mosquito production in cisterns and wells in St. Augustine, FL using attractive sugar baits. Acta Tropica. 2012; 124(2): 158–161.

20. Leta S, Beyene TJ, De Clercq EM, Amenu K, Kraemer MU, Revie CW. Global risk mapping for major diseases transmitted by *Aedes aegypti* and *Aedes albopictus*. International Journal of Infectious Diseases. 2018; 67: 25–35.

21. Ryan SJ, Carlson CJ, Mordecai EA, Johnson LR. Global expansion and redistribution of *Aedes*-borne virus transmission risk with climate change. PLoS Neglected Tropical Diseases. 2019; 13(3): e0007213.

22. Tian J, Mao G, Yu B, Fouad H, Wang C, Ga’al H, Mo J. Effect of common ornamental plants on the survivorship and fecundity of the *Aedes albopictus* (Diptera: Culicidae). Florida Entomologist. 2019; 102(1): 36–42.

23. Gouagna LC, Poueme RS, Dabiré KR, Ouédraogo JB, Fontenille D, Simard F. Patterns of sugar feeding and host plant preferences in adult males of *An. gambiae* (Diptera: Culicidae). Journal of Vector Ecology. 2010; 35(2): 267–276.

24. Martinez-Ibarra JA, Rodriguez MH, Arredondo-Jimenez JI, Yuval B. Influence of plant abundance on nectar feeding by *Aedes aegypti* (Diptera: Culicidae) in southern Mexico. Journal of Medical Entomology. 1997; 34(6): 589–593.

25. Li Y, Kamara F, Zhou G, Puthiyakunnon S, Li C, Liu Y, Zhou YL, Yao GY, Chen XG. Urbanization increases *Aedes albopictus* larval habitats and accelerates mosquito development and survivorship. PLoS Neglected Tropical Diseases. 2014; 8(11): e3301.

26. Fikrig K, Peck S, Deckerman P, Dang S, St Fleur K, Goldsmith H, Qu S, Rosenthal H, Harrington LC. Sugar feeding patterns of New York *Aedes albopictus* mosquitoes are affected by saturation deficit, flowers, and host seeking. PLoS Neglected Tropical Diseases. 2020; 14(10): e0008244.

27. Gillett JD, Haddow AJ, Corbet PS. The Sugar-Feeding-Cycle in a Cage-Population of Mosquitoes. Ent. Exp. & Appl. 1962; 5(3): 223–232.

28. Yee WL, Foster WA. Diel sugar-feeding and host-seeking rhythms in mosquitoes (Diptera: Culicidae) under laboratory conditions. Journal of Medical Entomology. 1992; 29(5): 784–791.

29. Van Handel E. Rapid determination of glycogen and sugars in mosquitoes. Journal of American Mosquito Control. 1985; 1(3): 299–301.

30. R Core Team. R Foundation for Statistical Computing. R: A Language and Environment for Statistical Computing. 2017; https://www.R-project.org/

31. Kline DL. Traps and trapping techniques for adult mosquito control. Journal of the American Mosquito Control Association. 2006; 22(3): 490–496.

32. Wanjiku CD, Tchouassi P, Sole CL, Pirk C, Torto B. Plant sugar feeding patterns of wild-caught *Aedes aegypti* from dengue endemic and non-endemic areas of Kenya. Medical and Veterinary Entomology. 2021; 35(3): 417–425.

33. Nyasembe VO, Tchouassi DP, Pirk CWW, Sole CL, Torto B. Host plant forensics and olfactory-based detection in Afro-tropical mosquito disease vectors. PLoS Neglected Tropical Diseases. 2018; 12(2): e0006185.

34. Knab F. Mosquitoes as flower visitors. Journal of the New York Entomological Society. 1907; 15(4): 215–219.

35. Sarma R, Adhikari K, Khanikor B. Evaluation of efficacy of pinene compounds as mosquitocidal agent against *Aedes aegypti* Linn.(Diptera: culicidae). International Journal of Tropical Insect Science. 2022; 42(3): 2567–2577.

36. Russo EB, Marcu J. Cannabis pharmacology: the usual suspects and a few promising leads. Advances in Pharmacology. 2017; 80: 67–134.

37. Liakakou A, Angelis A, Papachristos DP, Fokialakis N, Michaelakis A, Leandros A, Skaltsounis LA. Isolation of Volatile Compounds with Repellent Properties against *Aedes albopictus* (Diptera: Culicidae) Using CPC Technology. Molecules. 2021; 26(11): 3072.

38. Najar-Rodriguez AJ, Galizia CG, Stierle J, Dorn S. Behavioral and neurophysiological responses of an insect to changing ratios of constituents in host plant-derived volatile mixtures. Journal of Experimental Biology. 2010; 213(19): 3388–3397.

39. Tunón H, Thorsell W, Bohlin L. Mosquito repelling activity of compounds occurring in *Achillea millefolium* L.(asteraceae). Economic Botany. 1994; 48(2): 111–120.

40. Lee DC, Ahn YJ. Laboratory and simulated field bioassays to evaluate larvicidal activity of Pinus densiflora hydrodistillate, its constituents and structurally related compounds against *Aedes albopictus*, *Aedes aegypti* and *Culex pipiens pallens* in relation to their inhibitory effects on acetylcholinesterase activity. Insects. 2013; 4(2): 217–229.

41. Boonyuan W, Grieco JP, Bangs MJ, Prabaripai A, Tantakom S, Chareonviriyaphap T. Excito-repellency of essential oils against an *Aedes aegypti* (L.) field population in Thailand. Journal of Vector Ecology. 2014; 39(1): 112–122.

42. da Silva RCS, Milet-Pinheiro P, Bezerra da Silva PC, da Silva AG, da Silva MV, Navarro DMDAF, da Silva NH. (E)-caryophyllene and α-humulene: Aedes aegypti oviposition deterrents elucidated by gas chromatography-electrophysiological assay of Commiphora leptophloeos leaf oil. PLoS One. 2015; 10(12): e0144586.

43. Fenske MP, Hazelton KDH, Hempton AK, Shim JS, Yamamoto BM, Riffell JA, Imaizumi T. Circadian clock gene LATE ELONGATED HYPOCOTYL directly regulates the timing of floral scent emission in Petunia. Proceedings of the National Academy of Sciences. 2015; 112(31): 9775–9780.

44. Clements AN. The biology of mosquitoes. Volume 2: Sensory reception and behaviour. CABI publishing; 1999.

45. Dube FF, Tadesse K, Birgersson G, Seyoum E, Tekie H, Ignell R, Hill SR. Fresh, dried or smoked? Repellent properties of volatiles emitted from ethnomedicinal plant leaves against malaria and yellow fever vectors in Ethiopia. Malaria Journal. 2011; 10(1): 375.

46. Tucker KR, Steele CH, McDermott EG. *Aedes aegypti* (L.) and *Anopheles stephensi* Liston (Diptera: Culicidae) Susceptibility and Response to Different Experimental Formulations of a Sodium Ascorbate Toxic Sugar Bait. Journal of Medical Entomology. 2022; 59(5): 1710–1720.

47. Davis TJ, Kline DL, Kaufman PE. *Aedes albopictus* (Diptera: Culicidae) oviposition preference as influenced by container size and *Buddleja davidii* plants. Journal of Medical Entomology. 2016; 53(2): 273–278.

48. Cantrell CL, Ali A, Jones AMP. Isolation and identification of mosquito biting deterrents from the North American mosquito repelling folk remedy plant, *Matricaria discoidea* DC. PloS One. 2018; 13(10).

49. Wheelwright M, Catherine RW, Riabinina O. Olfactory systems across mosquito species. Cell and Tissue Research. 2021; 383(1): 75–90.

50. Farré-Armengol G, Filella I, Llusià J, Peñuelas J. β-Ocimene, a key floral and foliar volatile involved in multiple interactions between plants and other organisms. Molecules. 2017; 22(7): 1148.

51. Potter DA, Redmond CT, McNamara TD, Munshaw GC. Dwarf White Clover Supports Pollinators, Augments Nitrogen in Clover–Turfgrass Lawns, and Suppresses Root-Feeding Grubs in Monoculture but Not in Mixed Swards. Sustainability. 2021; 13(21): 11801.

52. Jakobsen HB, Kritjánsson K. Influence of temperature and floret age on nectar secretion in *Trifolium repens* L. Annals of Botany. 1994; 74(4): 327–334.

53. Kicel A, Wolbis M, Kalemba D, Wajs A. Identification of volatile constituents in flowers and leaves of *Trifolium repens* L. Journal of Essential Oil Research. 2010; 22(6): 624–627.

54. Xu P, Zhu F, Buss GK, Leal WS. 1-Octen-3-ol - the attractant that repels. F1000. Research. 2015; 4.

55. Shone SM, Ferrao PN, Lesser CR, Glass GE, Norris DE. Evaluation of Carbon Dioxide– and 1-Octen-3-Ol–Baited Centers for Disease Control Fay–Prince Traps to Collect *Aedes Albopictus*. Journal of the American Mosquito Control Association. 2003; 19(4): 445.

56. Upchurch RP, Peterson ML, Hagan RM. Effect of Soil-Moisture Content on the Rate of Photosynthesis and Respiration in Ladino Clover (*Trifolium Repens* L.). Plant Physiology. 1955; 30(4): 297.

57. Gibson PB, Cope WA. White Clover. Clover Science and Technology. 1985; 25: 471–490.

58. Seebens H, Blackburn TM, Dyer EE, Genovesi P, Hulme PE, Jeschke JM, … Essl F. No saturation in the accumulation of alien species worldwide. Nature Communications. 2017; 8(1): 1–9.

59. Korányi D, Markó V. Host plant identity and condition shape phytophagous insect communities on urban maple (*Acer spp.*) trees. Arthropod-Plant Interactions. 2022; 16(1): 129–143.

60. Stone CM, Foster WA. Plant-sugar feeding and vectorial capacity. In Ecology of parasite-vector interactions. Wageningen Academic Publishers, Wageningen; 2013. 35–79.

61. Nasci RS. *Toxorhynchites rutilus septentrionalis* feeding on tree sap. Journal of the American Mosquito Control Association. 1986; 2(4).

62. Swammerdam J. The book of nature, or, the history of insects: reduced to distinct classes, confirmed by particular instances. 1758.

63. Peach DA, Gries G. Mosquito phytophagy–sources exploited, ecological function, and evolutionary transition to haematophagy. Entomologia Experimentalis et Applicata. 2020; 168(2): 120–136.

64. Wondwosen B, Birgersson G, Tekie H, Torto B, Ignell R, Hill SR. Sweet attraction: sugarcane pollen-associated volatiles attract gravid *Anopheles arabiensis*. Malaria Journal. 2018; 17(1): 1–9.

65. Cabelin VLD, Alejandro GJD. Efficiency of matK, rbcL, trnH-psbA, and trnL-F (cpDNA) to molecularly authenticate Philippine ethnomedicinal Apocynaceae through DNA barcoding. Pharmacognosy Magazine. 2016; 12: S384.

66. Liu Z, Zeng X, Yang D, Chu G, Yuan Z, Chen S. Applying DNA barcodes for identification of plant species in the family Araliaceae. Gene. 2012; 499(1): 76–80.

67. Loera-Sánchez M, Studer B, Kölliker R. DNA barcode trnH-psbA is a promising candidate for efficient identification of forage legumes and grasses. BMC research notes, 2020; 13(1): 1–6.

68. Pang X, Liu C, Shi L, Liu R, Liang D, Li H, … Chen S. Utility of the trnH–psbA intergenic spacer region and its combinations as plant DNA barcodes: a meta-analysis. PloS One. 2012; 7(11): e48833.

69. Pang X, Song J, Zhu Y, Xu H, Huang L, Chen S. Applying plant DNA barcodes for Rosaceae species identification. Cladistics. 20122; 27(2): 165–170.

70. Chalcoff VR, Aizen MA, Galetto L. Nectar concentration and composition of 26 species from the temperate forest of South America. Annals of Botany. 2006; 97(3): 413–421.

71. Peach DA, Gries G. Nectar thieves or invited pollinators? A case study of tansy flowers and common house mosquitoes. Arthropod-Plant Interactions. 2016; 10(6): 497–506.

